# Epilepsy-associated Variants of a Single SCN1A Codon exhibit Divergent Functional Properties

**DOI:** 10.1101/2025.10.16.682932

**Authors:** Lanie N. Liebovitz, Christopher H. Thompson, Linda L. Laux, Alfred L. George

**Affiliations:** Department of Pharmacology, Northwestern University Feinberg School of Medicine, Chicago IL USA; Ann and Robert H Lurie Children’s Hospital of Chicago, Chicago IL USA

**Author notes:** Correspondence should be addressed to Alfred L. George, Jr.

## Abstract

**Objective:** Pathogenic variants in SCN1A, which encodes the voltage gated sodium channel Na_V_1.1, are associated with multiple epilepsy syndromes exhibiting a range of clinical severity. Loss or gain of function SCN1A variants are reported in different syndromes including Dravet syndrome, which is associated with loss-of-function whereas neonatal/infantile-onset developmental and epileptic encephalopathy (DEE) is associated with gain-of-function. Strategies to predict SCN1A variant pathogenicity and dysfunction have been proposed but are limited by available training data. We investigated the functional properties of four epilepsy-associated SCN1A variants affecting the same codon and sought to correlate channel dysfunction with phenotype.

**Methods:** Whole-cell manual patch-clamp recording was performed on heterologously-expressed Na_V_1.1 variants. Structural modeling of Na_V_1.1 variant proteins was conducted using AlphaFold 3.

**Results:** We describe an individual with early infantile onset DEE associated with SCN1A-I1347T, and identified three additional cases from the literature or ClinVar with distinct variation of the same codon (I1347N, I1347V, I1347F). Functional studies demonstrated mixed gain and loss of function properties for I1347T, I1347V, and I1347F, but complete loss-of-function for I1347N. Structural models suggest important interactions between isoleucine-1347 and the sixth transmembrane helices of domains 3 and 4 that are disrupted most significantly with asparagine replacement at this position (I1347N).

**Interpretation:** Pathogenic variants in SCN1A involving the same codon can produce divergent functional effects. Our findings suggest that predicting specific functional effects of SCN1A variants should not rely heavily on position in the protein.

## Introduction

Genetic testing has a high diagnostic yield in infant and ne-onatal-onset idiopathic epilepsy.^1-4^ Among the most frequent genetic etiologies discovered are pathogenic variants in SCN1A, which encodes the voltage-gated sodium (Na_V_) channel Na_V_1.1 expressed predominantly in GABAergic interneurons. Pathogenic SCN1A variants cause a clinical spectrum ranging from genetic epilepsy with febrile seizures plus (GEFS+) to severe developmental and epileptic encephalopathies (DEE) including Dravet syndrome.^5-8^

The molecular pathogenesis of SCN1A-related disorders investigated by in vitro assessments of Na_V_1.1 dysfunction has revealed that pathogenic variants associated with Dravet syndrome and GEFS+ impair function (e.g., loss-of-function; LoF)^9,10^ whereas a smaller subset of variants associated with early-onset DEE exhibit functional properties reported as gain-of-function (GoF).^11^ Discriminating between these extremes of channel dysfunction has therapeutic implications. Specifically, LoF SCN1A variants are considered contraindications for Na_V_ channel blocking anti-seizure medications,^12^ whereas GoF variants are associated with clinical phenotypes that may respond to this class of medicines.^11^ Assessing the pathogenicity of epilepsy-associated ion channel gene variants using the guidelines proposed by the American College of Medical Genetics^13^ does not discriminate between distinct functional subsets of SCN1A variants. Furthermore, the large number of identified SCN1A variants precludes detailed functional studies of all variants. Various strategies have been described for predicting the functional consequences of *SCN1A* variants in the absence of experimental evidence.^14-16^ These predictive models are trained with extent data including empirical evidence from voltage-clamp recording experiments. Because there are limited numbers of variants for which functional evidence is known, the training of predictive models is an ongoing process. One strategy used to predict functional effects of *SCN1A* variants relies upon comparisons with known function of paralogous Na_V_ channel variants.^15^ This approach is based on the inherent assumption that mutation of the same codon in the same or paralogous gene will function similarly. There is limited evidence that this assumption is correct in all cases.

In this study, we investigated the functional properties of four *SCN1A* variants affecting the same codon that are associated with DEE. We observed a wide spectrum of channel dysfunction among the variants including complete LoF, predominant GoF and mixed functional properties. Our findings suggest that caution should be exercised when predicting specific functional effects of *SCN1A* variants based largely on position in the protein.

## Results

### Case Presentation

A six-and-a-half-year-old child was evaluated at Lurie Children’s Hospital of Chicago for a complex seizure disorder. The child was born following an uncomplicated planned Cesarean section at term to healthy parents with no family history of seizures or sudden unexplained death. Initial development was normal. At age 3 months, the child had an unprovoked 20–30-minute generalized convulsion upon awakening from sleep, and then a second seizure with status epilepticus 4 days later. By parent report, EEG was normal at this time. At age 7 months, hemiconvulsions and alternating hemiconvulsions began. At age 11 months, focal seizures with impaired awareness occurred that were sometimes prolonged. EEG at age 15 months demonstrated multifocal sharp waves, slow and disorganized background, but no concurrent clinical events. Myoclonic seizures were noted at age 2 years. Subsequent EEGs at age 3 years and older showed slow, disorganized generalized periodic epileptiform discharges, one absence seizure, and near continuous diffuse interictal epileptiform discharges during sleep. A diagnosis of electrical status epilepticus during sleep was made. In addition to seizures, the child was nonverbal, nonambulatory, and could not sit without assistance. There was truncal hypotonia, appendicular hypertonia and hyperkinetic movements including dystonia, choreoathetosis and myoclonus.

Genetic testing revealed a de novo heterozygous *SCN1A* variant (c.4040T>C (p.Ile1347Thr; I1347T) that was classified as pathogenic and was absent in gnomAD (v.4.1.0). This variant was reported previously in a large case series of early infantile epileptic encephalopathy with no specific clinical details.^23^ Another missense variant affecting the same codon (I1347V) and inferred to be GoF was reported in association with neonatal onset DEE.^11^ Two additional missense variants at this position are registered in ClinVar (I1347N, I1347F).^24^ The affected residue is located in the short intracellular linker between segments S4 and S5 in domain 3. All variants are predicted to be GoF with probabilities ranging from 0.76 (I1347N) to 0.82 (all others) using the funNCion tool.^14^

### Functional properties of *SCN1A*-I1347T

To determine the functional consequences of the *SCN1A*-I1347T variant, we conducted whole-cell voltage clamp recordings of HEK293T cells transiently transfected with WT or mutant Na_V_1.1 channels. To ascertain the spectrum of Na_V_1.1 dysfunction we measured multiple biophysical properties including peak current density, voltage-dependence of activation and inactivation, recovery from inactivation, frequency-dependent channel rundown, kinetics of fast inactivation, persistent sodium current and responses to a slow voltage ramp depolarization (Supplemental Fig. S1). Experiments with cells expressing either WT or mutant Na_V_1.1 were conducted in parallel and data from 4 separate transfections were pooled.

Averaged whole-cell current traces illustrate smaller overall amplitudes for I1347T channels compared with WT Na_V_1.1 (Fig. 1A). Compared with WT channels, the peak current density for I1347T was significantly smaller and shifted approximately 10 mV in the hyperpolarized direction (Fig. 1B). The time course of fast inactivation appeared slower for the mutant (Fig. 1C) and inactivation time constants were significantly larger at 10 and 20 mV (Fig. 1D). Currents elicited by a slow depolarizing voltage ramp from −60 to 0 mV were larger (Fig. 1E, Fig. 2A), and exhibited greater levels of persistent current (Fig. 2B). These findings indicate that I1347T impairs fast inactivation leading to enhanced sodium ion conductance over time.

**Fig. 1.**
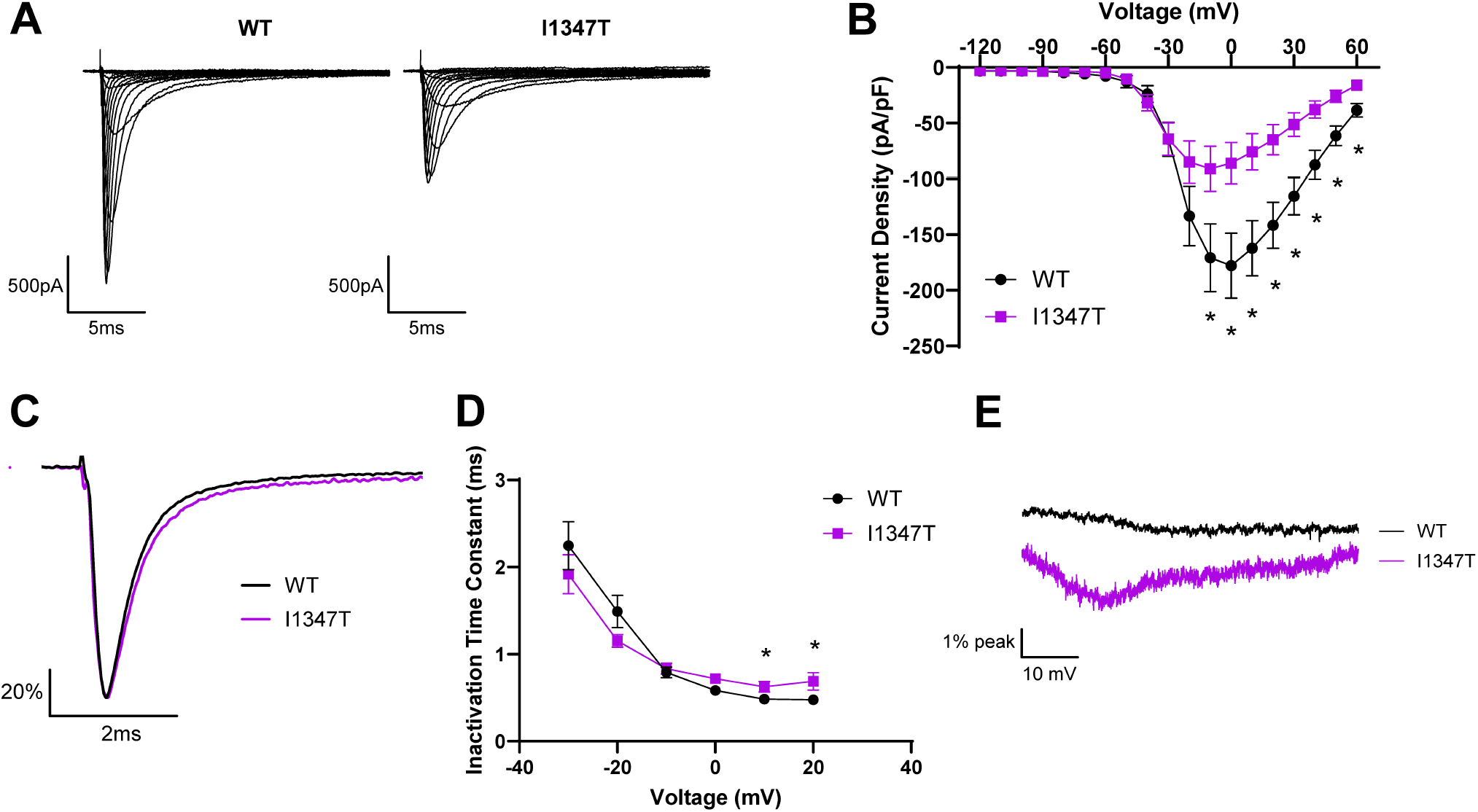
*SCN1A*-I1347T alters Na_V_1.1 functional properties. (A) Average whole-cell sodium current traces recorded from HEK293T cells expressing WT Na_V_1.1 or I1347T. (B) Current density-voltage relationship of WT Na_V_1.1 (n = 18) and I1347T (n = 20). *, *P* < 0.05 compared with WT. (C) Average current traces recorded at 0 mV and normalized to peak current to highlight differences in inactivation time-course exhibited by WT Na_V_1.1 and I1347T. (D) Voltage dependence of inactivation time constants determined for WT (n = 15) and I1347T (n = 15). *, *P* < 0.05 compared with WT. (E) Average ramp currents normalized to peak current from WT and I1347T. Data are expressed as mean ± SEM. Statistical comparisons were made using an unpaired Student’s *t* test. Complete quantitative data are summarized in **Supplemental Table S2**.

**Fig. 2.**
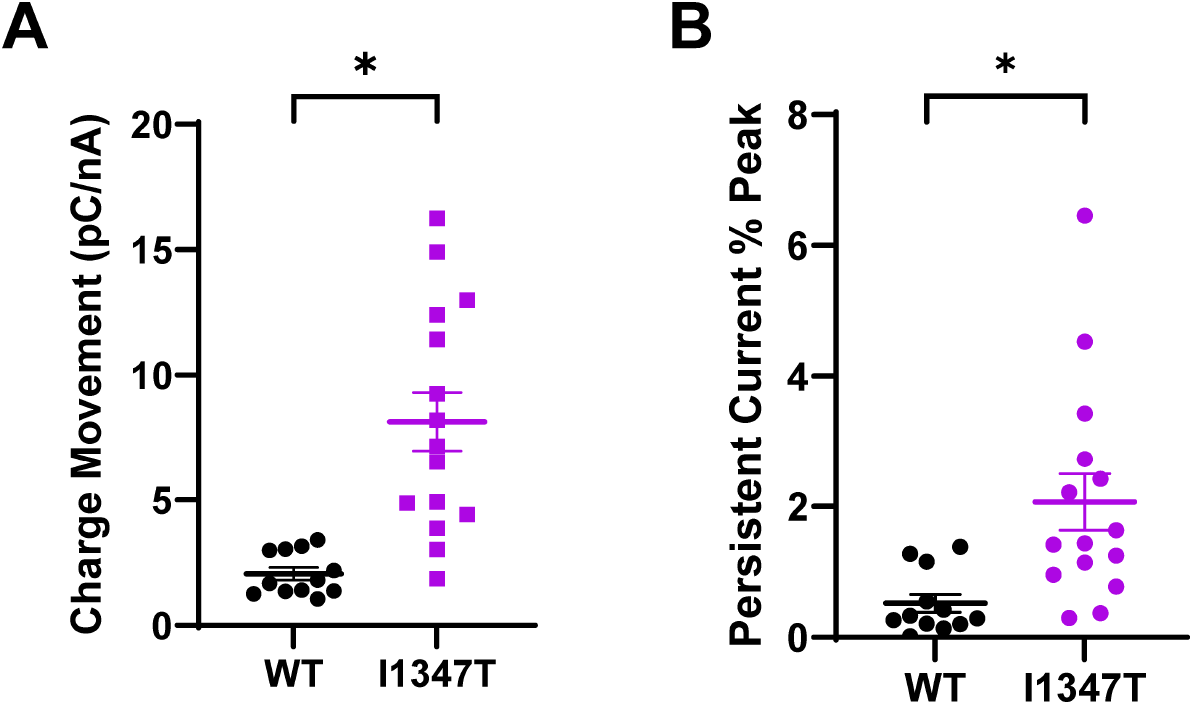
Ramp and persistent current for WT Na_V_1.1 and I1347T. (A) Charge movement elicited by a slow depolarizing voltage ramp recorded from HEK293T cells expressing WT Na_V_1.1 (n = 12) or I1347T (n = 15). (B) Persistent current exhibited by WT Na_V_1.1 (n = 12) and I1347T (n = 15) recorded during the ramp protocol. *, *P* < 0.05 compared with WT. Data are expressed as mean ± SEM. Statistical comparisons were made using an unpaired Student’s *t* test. Complete quantitative data are summarized in **Supplemental Table S2**.

The voltage dependence of activation and inactivation for I1347T channels were both hyperpolarized relative to WT Na_V_1.1 (Fig. 3A, Supplemental Table S2). The hyperpolarized shift in activation V½ observed for the mutant was larger than that for steady-state inactivation creating a large area of overlap between the two curves (‘window current’), which represents the range of membrane voltages where channels are activated but not inactivated (Fig. 3B). These differences in voltage-dependence along with enhanced window current suggest a net GoF. Lastly, we examined the kinetics of recovery from fast inactivation and the response to repetitive depolarizing pulsed to elicit use-dependent rundown. The time course of recovery from inactivation for I1347T was significantly slower than WT Na_V_1.1 (Fig. 3C) and this correlated with enhanced use-dependent rundown of the current in cells expressing the mutant channel (Fig. 3D). The slower recovery kinetics observed for I1347T is explained by larger time constants for both fast and slow components of recovery and a smaller fraction recovering with the fast component (Supplemental Table S2). Slower recovery from inactivation and enhanced use-dependent rundown are predicted to cause lower overall function during high frequency action potential firing. Overall, our functional analysis demonstrates that I1347T exhibits both GoF and LoF effects. This variant can be classified as having mixed dysfunction with some prominent GoF features.

**Fig. 3.**
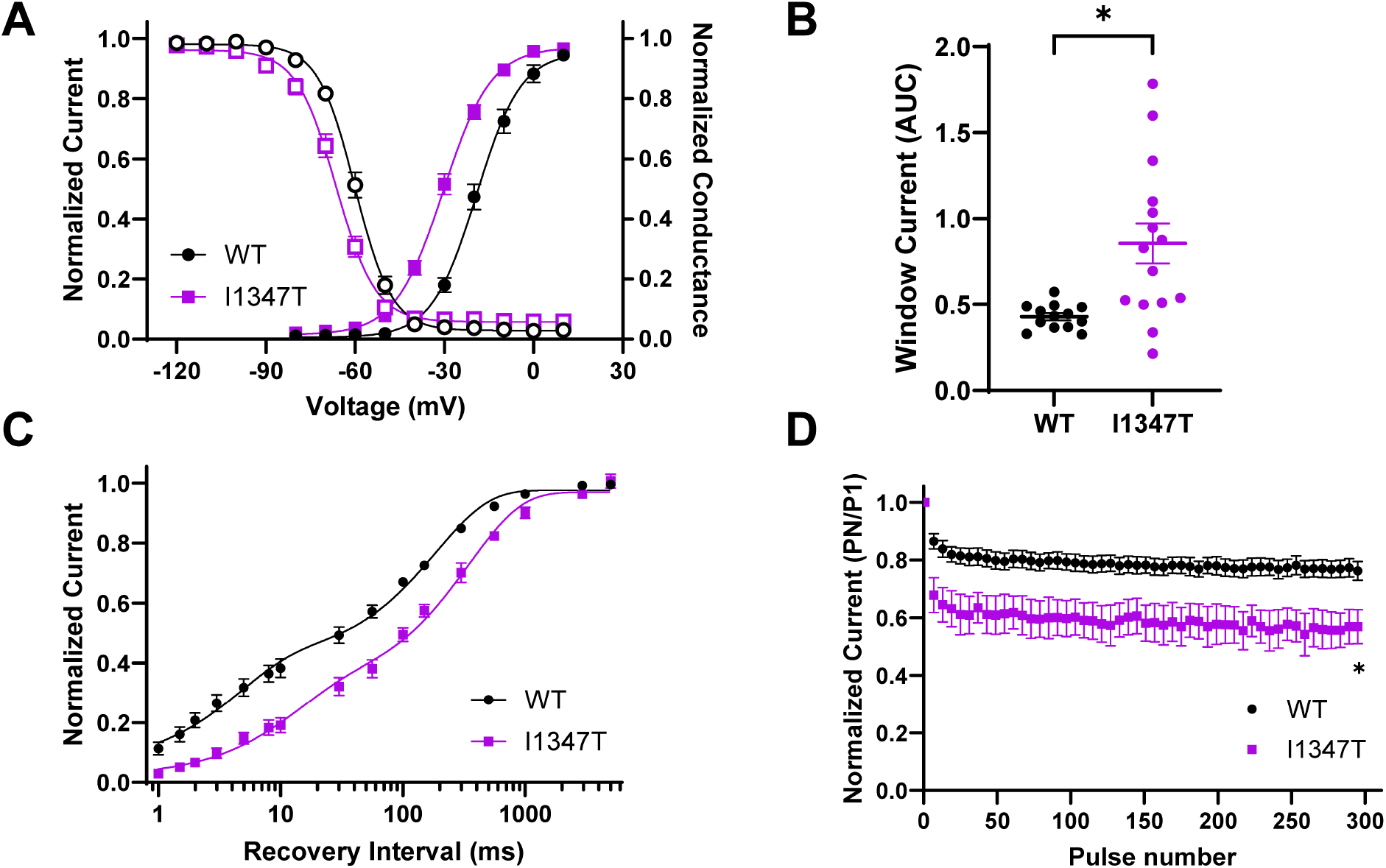
*SCN1A*-I1347T alters Na_V_1.1 voltage-dependence and recovery from inactivation. (A) Voltage dependence of activation overlayed with voltage dependence of inactivation of WT Na_V_1.1 (n = 15) and I1347T (n = 16). (B) Window current area quantified for WT Na_V_1.1 and I1347T. *, *P* < 0.05 compared with WT. (C) Time course of recovery from inactivation after 100 msec depolarization comparing WT Na_V_1.1 (n = 13) and I1347T (n = 14). (D) Plot of residual current after repetitive pulsing to 0 mV for WT Na_V_1.1 (n = 13) and I1347T (n = 15). *, *P* < 0.05 compared with WT. Data are expressed as mean ± SEM. Statistical comparisons were made using an unpaired Student’s *t* test. Complete quantitative data are summarized in **Supplemental Table S2**.

### Functional properties of other *SCN1A*-I1347 Variants

Three other missense variants involving *SCN1A* codon 1347 are reported in ClinVar (I1347N, I1347F) or in a published case series (I1347V; case 13 in 11. Phenotype information is limited for the two variants in ClinVar (DEE reported for I1347N; neonatal DEE reported for I1347F), but detailed clinical features associated with de novo I1347V were reported in a case series of *SCN1A* GoF variants. The reported individual with I1347V had tonic seizures starting day 3 of life evolving to intractable generalized tonic-clonic seizures with myoclonus, developmental regression after age 1 year with profound intellectual disability (non-verbal, non-ambulatory), generalized spike-wave pattern and continuous spike-wave during sleep by EEG, along with hyperkinesia and congenital arthrogryposis affecting all limbs.^11^ *SCN1A*-I1347V was inferred to be GoF based on an electrophysiological analysis of the equivalent mutation in SCN8A (I1327V), which is a recurrent pathogenic DEE-associated variant.^25-27^ There were no other paralogous variants with known functional properties in the SCN viewer database.^15^ We determined the functional consequences of these additional *SCN1A* I1347 variants and compared findings to WT channels assayed in parallel.

Heterologous expression of *SCN1A*-I1347N in HEK293T cells generated sodium current that was not significantly different from non-transfected cells (Fig. 4). The same observation was made with cells transfected with a second independent recombinant plasmid clone of this mutation. In total, data were analyzed from 6 separate transfections. The level of current density precluded determination of other biophysical properties, and we concluded that *SCN1A*-I1347N is a complete LoF variant.

**Fig. 4.**
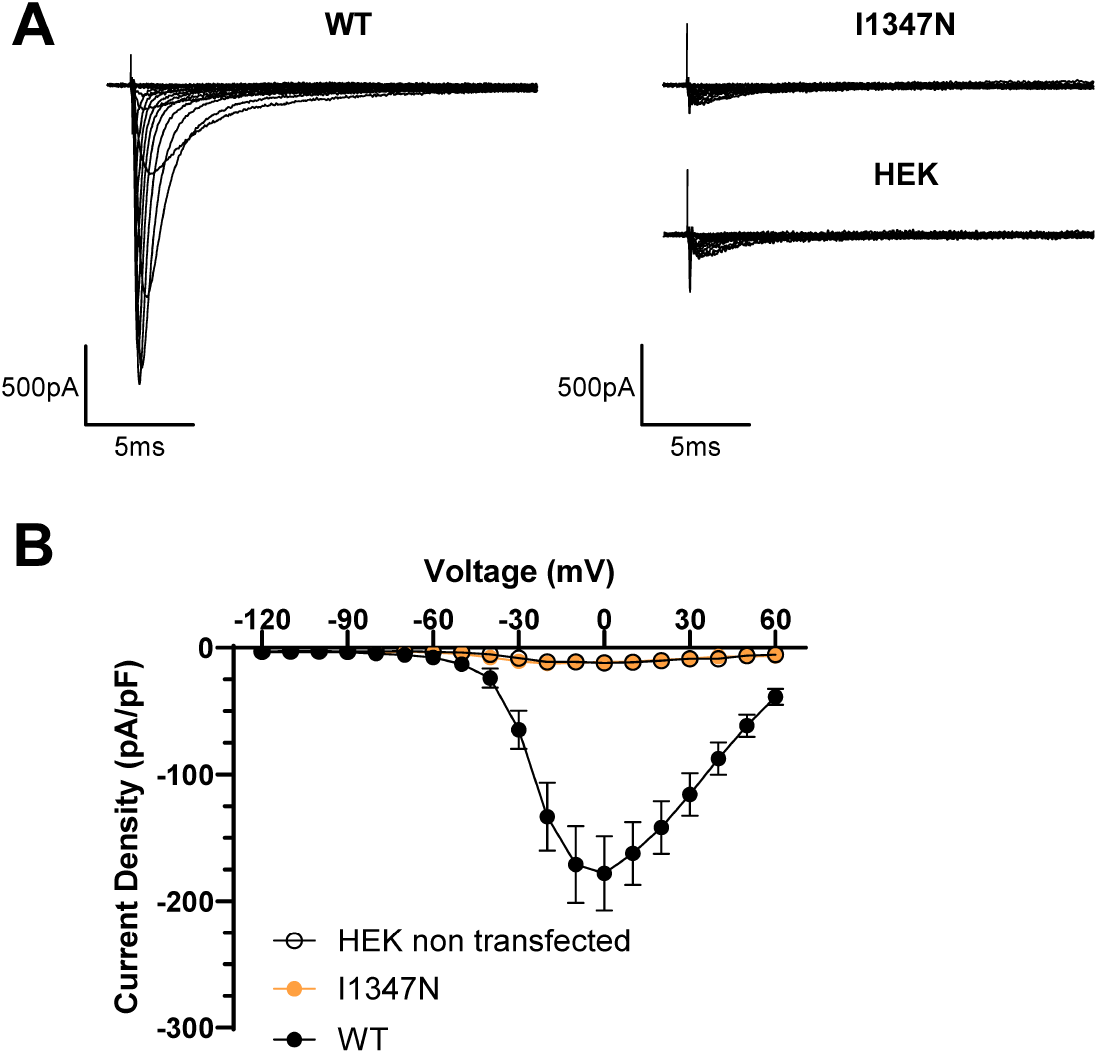
*SCN1A*-I1347N encodes a non-functional Na_V_1.1 channel. (A) Average whole-cell sodium current traces from HEK293T cells expression WT Na_V_1.1 or I1347N, compared with non-transfected cells. (B) Current density-voltage relationship of WT Na_V_1.1 (n = 18), I1347N (n = 16), and non-transfected HEK cells (n = 9). Complete quantitative data are summarized in **Supplemental Table S2**.

Transient expression of *SCN1A*-I1347V or *SCN1A*-I1347F in HEK293T cells yielded measurable sodium currents that were compared to cells expressing WT channels. Data from 2 separate transfections were pooled for I1347V, and data from 3 separate transfections were pooled for I1347F. The WT data measured in parallel with I1347V and I1347F were pooled to perform statistical comparisons with the two Na_V_1.1 variants.

Averaged whole-cell current traces illustrate smaller overall amplitudes for I1347V and I1347F channels compared with WT Na_V_1.1 (Fig. 5A). The peak current densities for I1347V and I1347F overlapped and were significantly smaller compared with WT channels (Fig. 5B). The time course of fast inactivation was slower for I1347F and I1347V in comparison to WT channels (Fig. 5C). Correspondingly, inactivation time constants were significantly larger than WT for I1347F and I1347V at 0 mV (Fig. 5D, Table S2). Currents elicited by a slow depolarizing voltage ramp from −60 to 0 mV were larger than WT channels for both Na_V_1.1 variants, with I1347F having the greatest difference from WT (Fig. 5E, Fig. 6A). I1347F also exhibited a greater level of persistent current than WT and I1347V channels (Fig. 6B). The time course of recovery from inactivation for I1347V appears slower than WT Na_V_1.1 (Fig. 7A) and this correlated with enhanced use-dependent rundown of the current in cells expressing the I1347V mutant channel (Fig. 7B). The time course of recovery from inactivation for I1347F appears faster than WT Na_V_1.1 and mutant I1347V (Fig. 7A), while the use-dependent rundown of current in cells from I1347F was not different from WT Na_V_1.1 (Fig. 7B). The faster recovery kinetics observed for I1347F is explained by a larger fraction recovering with the fast component (Supplemental Table S2). The voltage dependence of activation and inactivation for I1347V or I1347F were not significantly different from WT Na_V_1.1 (Supplemental Fig. S2). Overall, our functional analysis demonstrates that I1347V and I1347F exhibit a mixture of GoF and LoF effects. Supplemental Table S3 presents a summary of the observed functional abnormalities we observed for all functional *SCN1A*-I1347 variants.

**Fig. 5.**
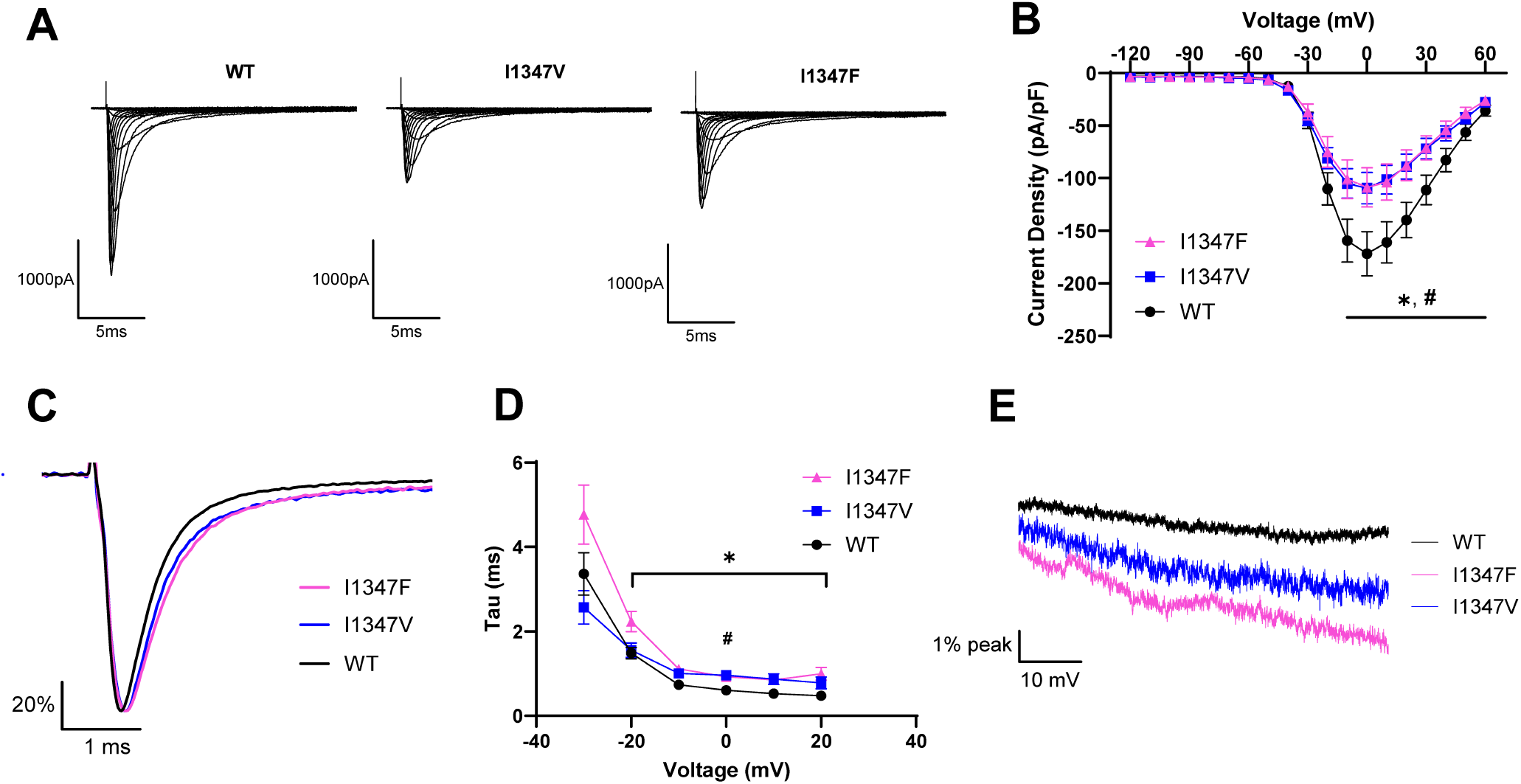
Functional properties of *SCN1A* variants I1347V and I1347F. (A) Average whole-cell sodium current traces recorded from HEK293T cells expressing WT Na_V_1.1, I1347V, or I1347F. (B) Current density-voltage relationship of WT Na_V_1.1 (n = 18), I1347V (n = 18), and I1347F (n = 17). *, *P* < 0.05 for I1347F compared with WT. #, *P* < 0.05 for I1347V compared with WT. (C) Average current recorded at 0 mV and normalized to peak current to highlight differences in inactivation time-course exhibited by WT Na_V_1.1, I1347V, and I1347F. (D) Voltage dependence of inactivation time constants determined for WT Na_V_1.1 (n = 18), I1347V (n = 18), and I1347F (n = 17). *, *P* < 0.05 for I1347F compared with WT. #, *P* < 0.05 for I1347V compared with WT. (E) Average ramp currents normalized to peak current from WT Na_V_1.1, I1347V, and I1347F. Data are expressed as mean ± SEM. Statistical comparisons were made using an unpaired Student’s *t* test. Complete quantitative data are summarized in **Supplemental Table S2**.

**Fig. 6.**
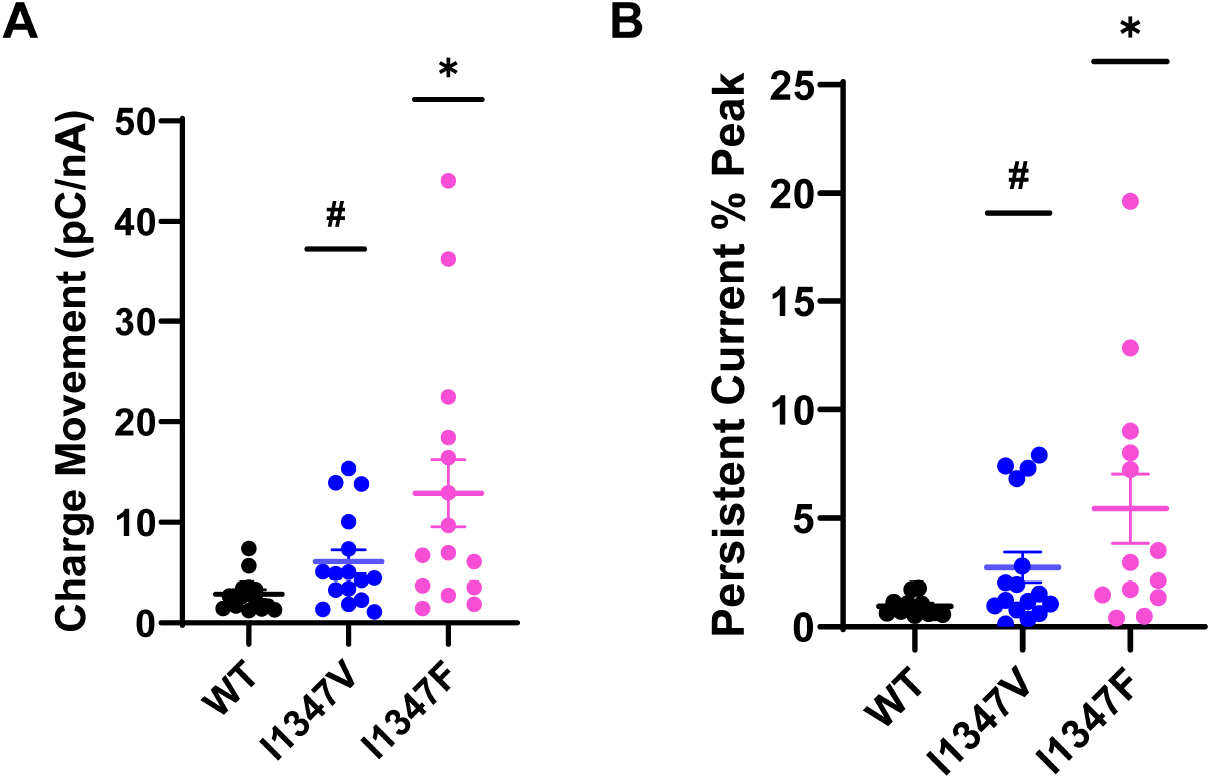
Ramp and persistent current for WT Na_V_1.1, I1347V, and I1347F. (A) Charge movement elicited by a slow depolarizing voltage ramp recorded from HEK293T cells expressing WT Na_V_1.1 (n = 14), I1347V (n = 16), and I1347F (n = 15). *, *P* < 0.05 for I1347F compared with WT. #, *P* < 0.05 for I1347V compared with WT. (B) Persistent current of WT Na_V_1.1 (n = 11), I1347V (n = 16), and I1347F (n = 13) recorded during the ramp protocol. *, *P* < 0.05 for I1347F compared with WT. #, *P* < 0.05 for I1347V compared with WT. Data are expressed as mean ± SEM. Statistical comparisons were made using an unpaired Student’s *t* test. Complete quantitative data are summarized in **Supplemental Table S2**.

**Fig. 7.**
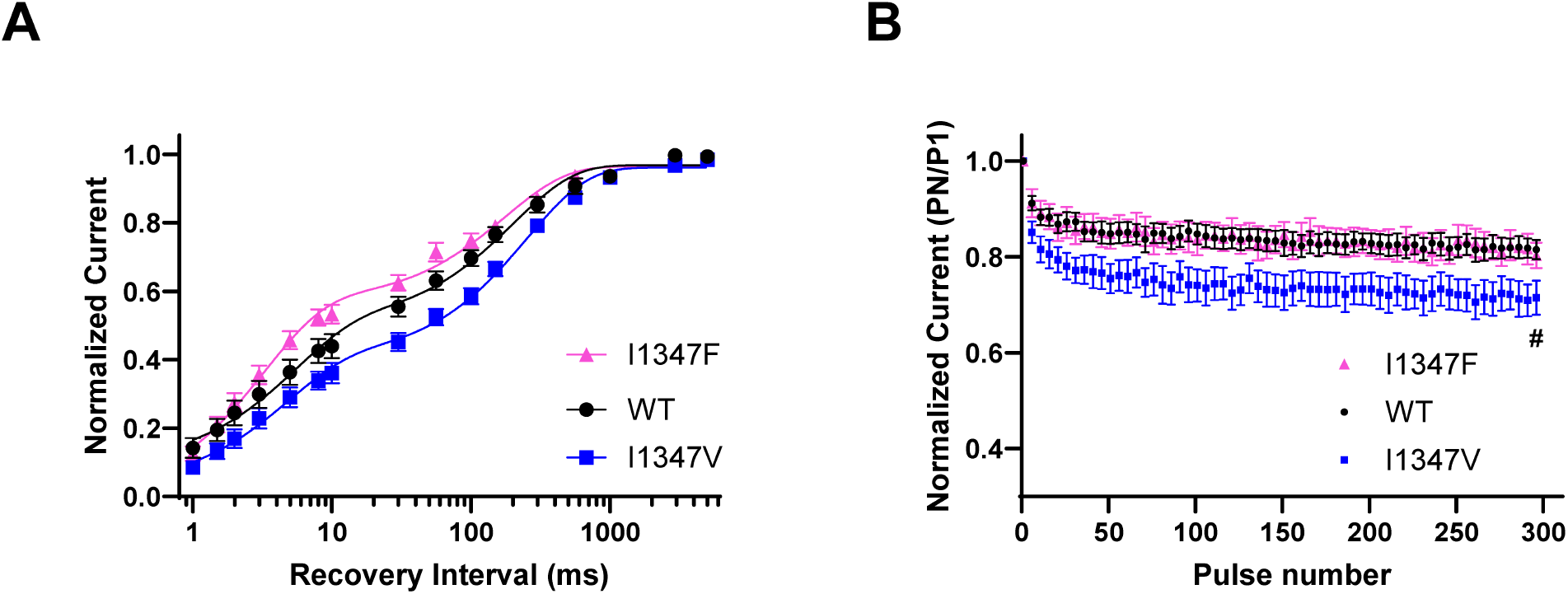
*SCN1A* variants I1347V and I1347F alter Na_V_1.1 inactivation recovery kinetics. (A) Time course of recovery from inactivation after 100 msec depolarization comparing WT Na_V_1.1 (n = 15), I1347V (n = 16), and I1347F (n = 16). (B) Plot of residual current after repetitive pulsing to 0 mV for WT Na_V_1.1 (n = 16), I1347V (n = 18), and I1347F (n = 16). #, *P* < 0.05 for I1347V compared with WT. Data are expressed as mean ± SEM. Statistical comparisons were made using an unpaired Student’s *t* test. Complete quantitative data are summarized in **Supplemental Table S2**.

### Structural basis for *SCN1A*-I1347 variant dysfunction

*SCN1A* variants at position 1347 in Na_V_1.1 exhibit a mixture of different GoF and LoF effects, with I1347N showing complete LoF. We constructed structural models of WT and each variant at position 1347, which is located at the junction of the domain 3 (D3) S4-S5 linker and the D3/S5 helix, to gain structural insights into the range of functional effects. In the WT channel, isoleucine-1347 interacts with the S6 helix of the same domain, as well as the S6 helix of domain 4 (D4/S6), specifically with leucine-1775 located at the distal end of D4/S6 (Fig 8A,B). This contact is lost in I1347N, which results in repositioning of D4/S6 and partially occluding the pathway for permeating ions (Supplemental Fig. S3; Supplemental Movie S1). This repositioning of D4/S6 was only identified in I1347N, which could explain why this variant uniquely causes complete LoF.

**Fig. 8.**
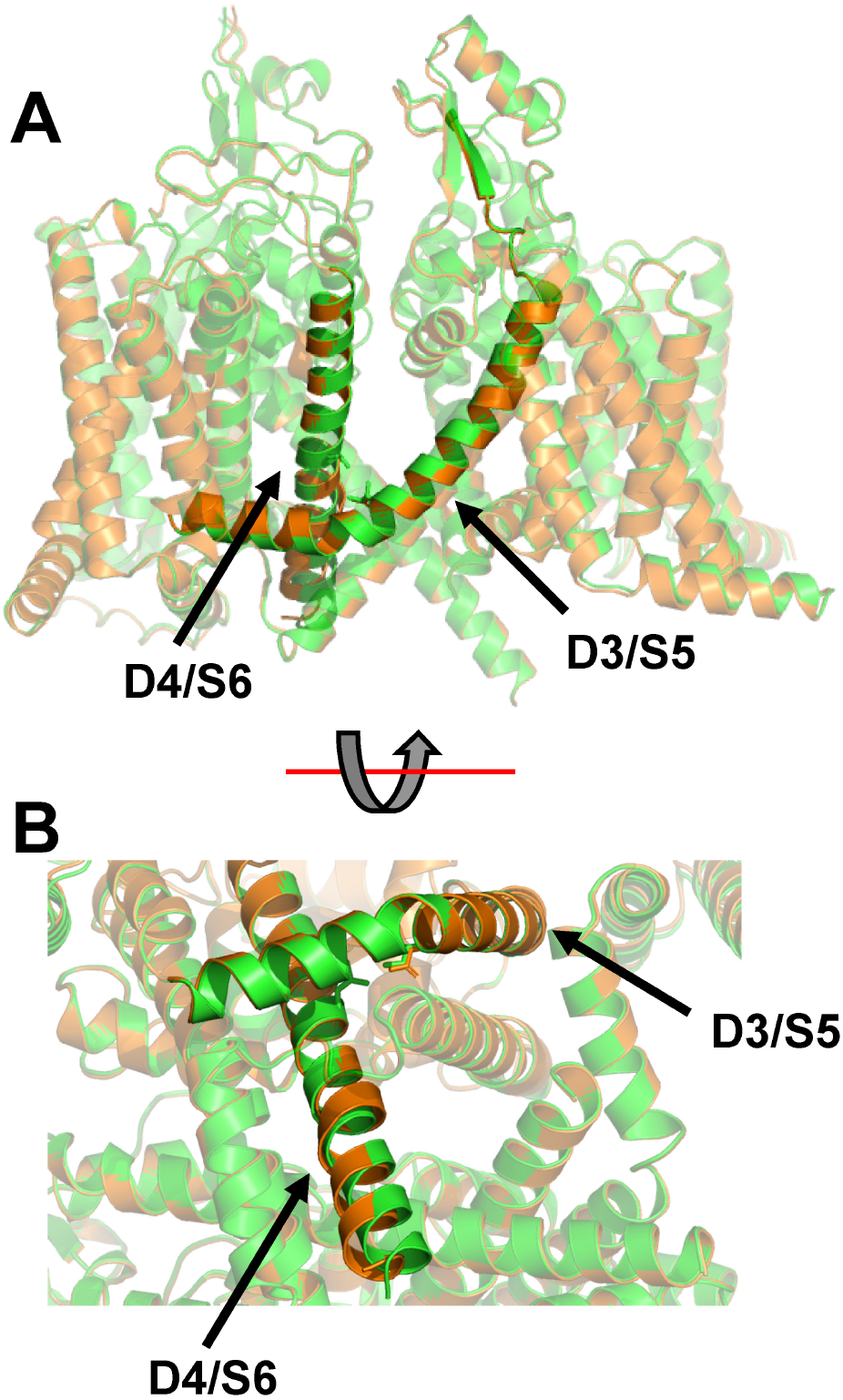
Structural modeling of Na_V_1.1-I1347N. (A) Structural alignment of Na_V_1.1-WT (green) and Na_V_1.1-I1347N (orange). (B) Zoomed and rotated view of segments D3/S5 and D4/S6 illustrating loss of interaction with of I1347 with L1775 and repositioning of segment D4/S6 in the mutant channel.

## Discussion

Dysfunctional brain Na_V_ channels including those encoded by *SCN1A*, SCN2A and SCN8A are well-known causes of neurodevelopmental disorders and various epilepsy syndromes._7_ Understanding the molecular pathogenesis of these conditions and delineating genotype-phenotype correlations contribute to improving diagnosis, predicting long-term outcomes, and designing precise therapy. Accurately classifying Na_V_ channel dysfunction helps predict the impact of variants on neuronal excitability and neural network activity, and for anticipating clinical responses to sodium channel blocking anti-seizure medications.

A major challenge encountered with increasing use of genetic testing is the rise in numbers of variants of uncertain significance (VUS). For channelopathies, conducting in vitro functional assessments of variants can provide information to help decrypt VUS, but performing this at scale with genetic variant discovery has not been feasible despite efforts to harness automation and mutagenesis screening._28-32_ For these reasons, computational prediction models have been developed as possible solutions to the VUS avalanche._14,33_ These models aspire to discriminate between disease-causing pathogenic variants and benign variants, but some also attempt to predict whether a variant confers GoF or LoF. Extensive training of such models is essential but there is limited empirical evidence of variant function. This necessitated extrapolating from what is known to interpolate functional properties of novel variants.

In this study, we investigated four *SCN1A* missense variants involving different amino acid substitutions of the same residue. We aligned the clinical phenotypes when known with in vitro functional properties of the associated variant. We described a case of a complex epilepsy syndrome and impaired neurodevelopment with early infantile onset associated with a *SCN1A* variant. This contributes to the emerging recognition of *SCN1A*-related early infantile DEE as a distinct clinical entity, which warrants different treatment considerations from the more common *SCN1A* LoF disorder Dravet syndrome. The I1347T variant exhibits mixed GoF and LoF properties, which may differ from other variants reported in this syndrome classified as simply GoF.

Our study illustrates challenges in extrapolating the functional impact of one variant to another even when located in the same protein location. Indeed, we illustrate that knowing the functional effects of I1347T (mixed GoF and LoF) would not have helped predict the complete LoF observed for a different amino acid substitution of the same residue (I1347N). Perhaps predictions based on extrapolations of paralogous variants should be interpreted with caution until more extensive data-driven training of the models can be accomplished. Intensifying efforts to experimentally determine functional properties of many more variants will help improve the accuracy of predictive models to classify channel dysfunction.

Our functional investigation of the previously reported variant I1347V inferred to be GoF also highlight the challenges in assigning simple functional labels to some variants. In that case, our experiments demonstrate that I1347V exhibits a mixture of GoF and LoF properties. The GoF properties stem largely from defects in fast inactivation, whereas the LoF features involve lower peak current density and slower recovery from inactivation. Applying selective interpretation rules (e.g., ignoring effects on current density and recovery from inactivation) would classify the variant as GoF. A similar classification would be made for I1347F, which also exhibits lower current density as a LoF feature. An unbiased systematic assessment of phenotype-function correlation in which functional properties are determined blind to clinical phenotype would be valuable.

Bridging the gap between biophysical studies and prediction of neural activity is the next challenge in neuronal channelopathies. Knowing which specific dysfunctional channel property or combination of defects drives the clinical phenotype requires more advanced studies involving human neurons, animal models or validated computational methods.

### Study Limitations

We used heterologous cells to investigate the functional properties of *SCN1A* variants and this approach may not be able to capture neuron-specific effects on trafficking and localization or that involve protein-protein interactions, which only occur in native cells. However, there are substantial challenges to studying individual mutations in native neurons including the presence of endogenous Na_V_ channels and the lack of Na_V_1.1 selective inhibitors to discriminate this channel from others. Introducing *SCN1A* mutations into induced pluripotent stem cells or genetically engineered mice are more advanced strategies that are beyond the scope of our study. Finally, two of the *SCN1A* variants we studied were from ClinVar and have limited phenotypic information making correlations between function and clinical features unreliable.

### Conclusion

We report a case with *SCN1A*-related early onset DEE associated with a mixed function variant and compared the functional properties of three other variants affecting the same amino acid position in the protein. Our findings demonstrate the challenges to predicting *SCN1A* variant function and offer additional insight into genotypephenotype relationships for this genetic disorder.

## Acknowledgements

The authors are grateful to Erin McGinnis for collecting clinical information about the index case. This work was supported by grant U54-NS108874 from the National Institute of Neurological Disease and Stroke and by the Baker Program in Undergraduate Research administered by Northwestern University Weinberg College of Arts and Sciences.

## Disclosures

Dr. George received research funding from Biohaven Pharmaceuticals for unrelated studies. Dr. George also consults with Tevard Biosciences, Praxis Precision Medicines and Neurocrine Biosciences for unrelated work. None of the other authors have any conflicts of interest to disclose. Dr. Laux has participated in research sponsored by GW Pharma, Zogenix, Biocodex, Stoke Therapeutics, Encoded Therapeutics, Epygenix therapeutics, Xenon, Praxis, Neurocrine, and Ovid Therapeutics.

## Materials and Methods

### Study Participants

A proband with a novel *SCN1A* variant was deidentified prior to inclusion in this research study. Approval for retrospective review of the case was obtained from the Institutional Review Board of the Ann and Robert H. Lurie Children’s Hospital of Chicago. All other *SCN1A* variants were obtained from the literature or ClinVar.

### Mutagenesis and heterologous expression of Na_V_1.1

Site-directed mutagenesis of recombinant intron-stabilized human Na_V_1.1 was performed using Q5 Hot Start High-Fidelity 2X Master Mix (New England Biolabs, Ipswich, MA) as described previously.^17^ Mutagenic primers were designed using custom software (Q5 primer designer; available at https://prism.northwestern.edu/records/tcsgw-z4295), and are presented in Supplemental Table S1. Full-length cDNA encoding WT or variant Na_V_1.1 was expressed from plasmids that also encoded an encephalomyocarditis virus internal ribosomal entry site with the wildtype A6 in the JK bifurcation loop^18^ followed by the monomeric red fluorescent protein mScarlet (pIRES2-mScarlet).

Complete sequence of each variant plasmid was performed with nanopore sequencing (Primordium Labs, Arcadia, CA) and analyzed using a custom multiple sequence alignment tool (MuSIC; available at https://prism.northwestern.edu/records/h6hc6-n0j20).

### Cell culture and transfection

HEK293T cells (CRL-3216; American Type Culture Collection, Manassas, VA) were transiently transfected with 2 µg WT or mutant Na_V_1.1 using PolyFect reagent (Qiagen, Germantown, MD). When a variant showed no activity, the transfection was repeated with another independent clone of the mutant. The HEK293T cell line was maintained in Dulbecco’s Modified Eagle Medium (GIBCO, Brooklyn, NY) with 10% fetal bovine serum (R&D Systems, Minneapolis, MN), 2 mM L-glutamine, 50 U/ml penicillin, and 50 µg/ml streptomycin at 37°C in 5% CO2. In some experiments, HEK293T cells were cultured overnight at 28°C to test effects of low temperature incubation on current density.

### Manual patch-clamp recording

Whole-cell voltage-clamp recordings of HEK293T cells were performed by conventional manual patch clamp as described previously.^19^ Recordings were performed at room temperature 24 and 48 hours after transfections. Data were acquired using an Axopatch 200B amplifier (Molecular Devices, San Jose, CA) at 50 kHz and filtered at 5 kHz. Recordings began 10 min after achieving the whole-cell configuration to allow for equilibration of pipette solution with intracellular fluid. Liquid junction potential was corrected at the beginning of each recording, series resistance was compensated 90%, and leak currents were subtracted using a P/4 procedure. Borosilicate glass capillaries (Harvard Apparatus, Holliston, MA) were used to fabricate patch pipettes using a multistage P-1000 Flaming-Brown micropipette puller (Sutter Instruments, Novato, CA) and were fire-polished using a microforge (MF-830; Narashige, Amityville, NY) to a resistance of 1.5–2.3 MΩ. The pipette solution consisted of (in mM) 10 NaF, 105 CsF, 20 CsCl, 2 EGTA, 10 HEPES, and 10 dextrose with the final pH adjusted to 7.35 with CsOH and osmolality adjusted to 300 mOsm/kg with sucrose. Bath (extracellular) solution containing (in mM) 140 NaCl, 4 KCl, 1.8 CaCl2, 1 MgCl2, 10 HEPES, and 10 dextrose, with the final pH adjusted to 7.35 with NaOH and osmolality adjusted to 310 mOsm/kg with sucrose.

### Data analysis

Data were analyzed and plotted with a combination of Clampfit 10.7 (Molecular Devices), Microsoft Excel (Microsoft, Redmond, WA), and GraphPad Prism (GraphPad Software, Boston, MA). Data are presented as mean ± SEM. Current traces presented in the figures are averages that were not corrected for cell capacitance. To illustrate the inactivation time course, currents recorded at the voltage that yielded the peak current were averaged and normalized to the maximum. Whole-cell currents were normalized to membrane capacitance to calculate current density. GraphPad Prism was used to fit voltage dependence of activation and inactivation curves with Boltzmann functions, time course of inactivation with a single exponential function, and recovery from inactivation with a two-exponential function. Window current was calculated by integrating the area under the intersection between the Boltzmann fits for voltage dependence of activation and inactivation using a custom MatLab script.^20^ All data were normalized to WT measured in parallel and were compared using Student’s t-test. The threshold for statistical significance was P ≤ 0.05. Exact P values are presented in the supplemental tables.

### Structural Analysis of Na_V_1.1

Structural models of Na_V_1.1 WT and variants were generated using AlphaFold 3.^21^ Five structures were generated and reviewed for Na_V_1.1 WT and each variant to reduce sampling bias. To ensure high confidence of the models, only regions of the protein with corresponding cryo-EM structures were included in the analysis, which corresponded to >80% confidence by the predicted local distance difference test.^22^ Hydrogen bonds and other side-chain interactions were identified using UCSF-Chimera. All visualization and structure figures were generated using PyMol 3.

**Fig. S1.**
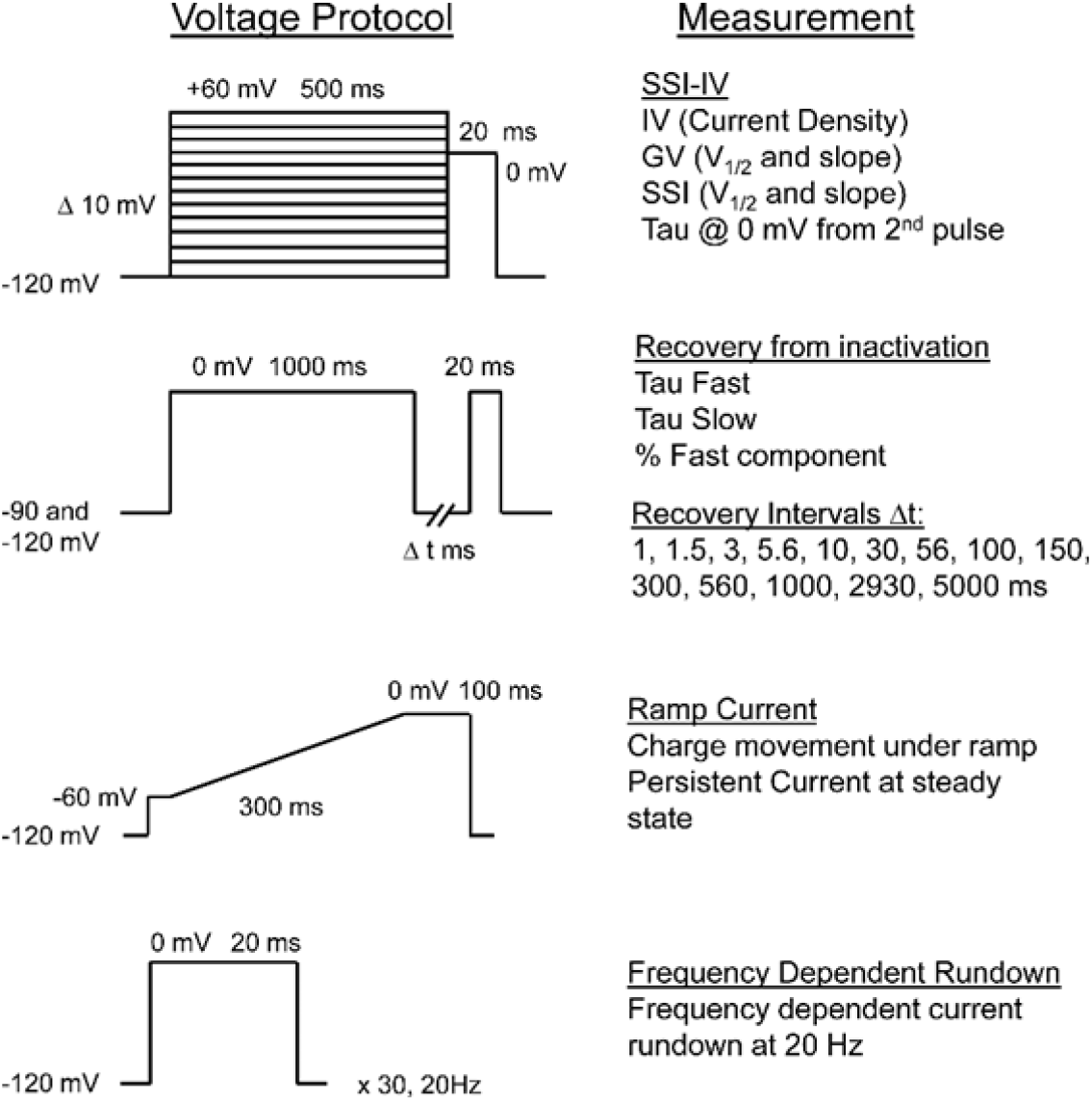
Voltage protocols used to assess biophysical parameters of Na_V_1.1.

**Fig. S2.**
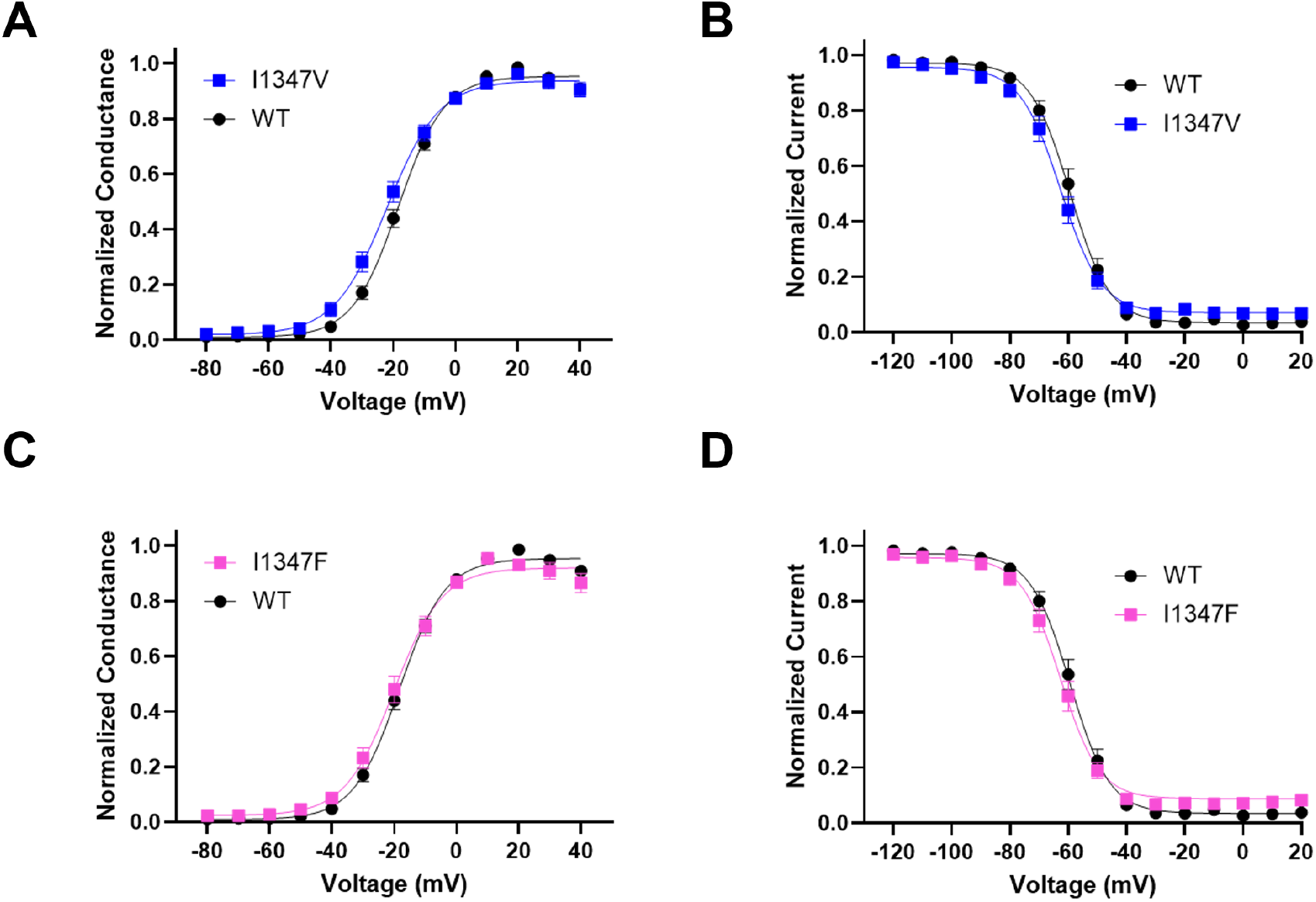
Effects of I1347V and I1347F on Na_V_1.1 voltage dependence of activation and inactivation. (A) Voltage dependence of channel activation of WT Na_V_1.1 (n = 18) and I1347V (n = 16). (B) Voltage dependence of channel inactivation of WT Na_V_1.1 (n = 18) and I1347V (n = 17). (C) Voltage dependence of channel activation of WT Na_V_1.1 (n = 18) and I1347F (n = 16). (D) Voltage dependence of channel inactivation of WT Na_V_1.1 (n = 18) and I1347F (n = 17). Data are expressed as mean ± SEM. Complete quantitative data are summarized in Supplemental Table S2 and Supplemental Dataset D1.

**Fig. S3.**
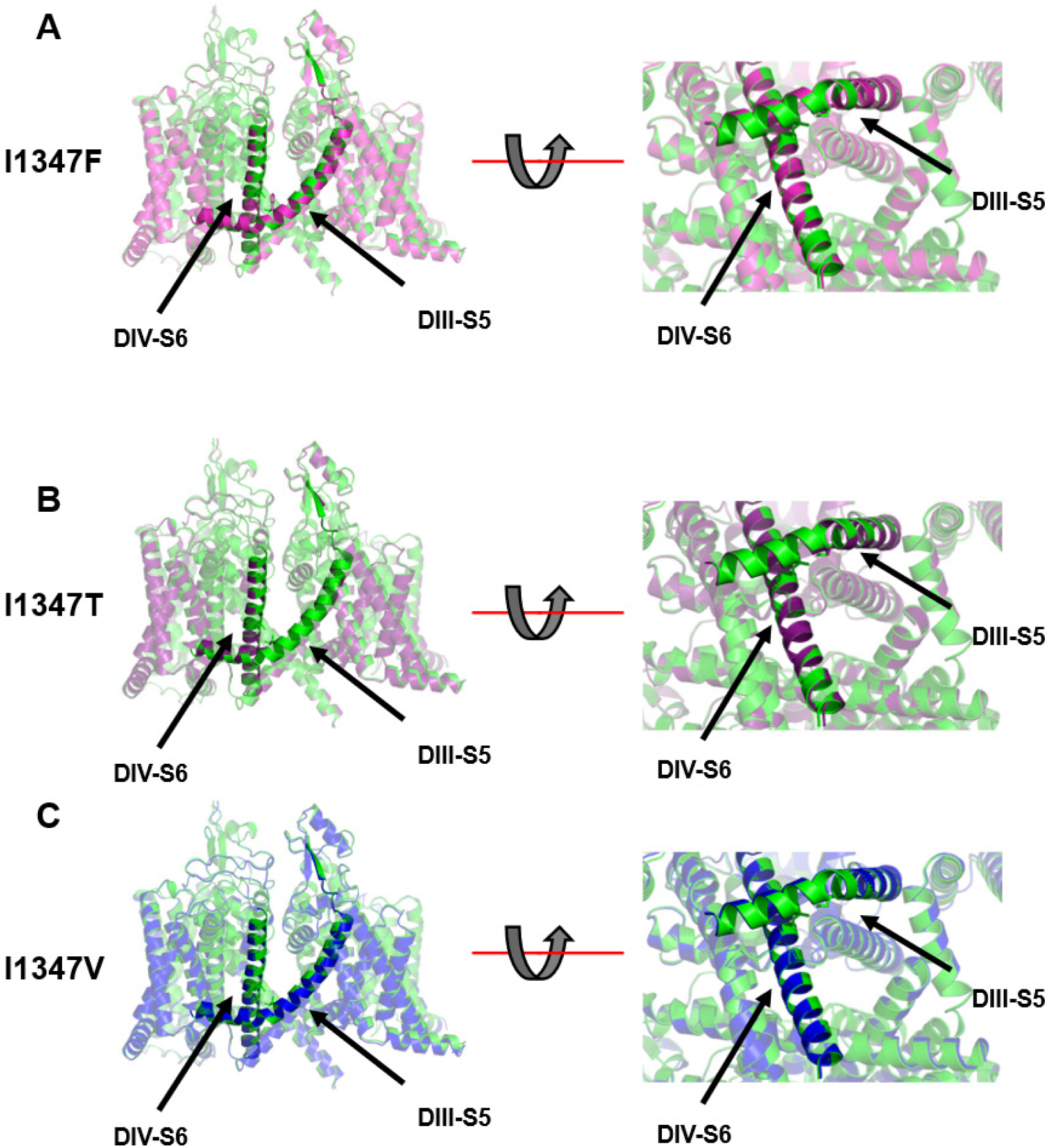
Structural analysis of variants at position 1347. Structural alignments of the full channel (left), and zoomed view of Domain III-S6 and Domain IV-S6 of (A) Na_V_1.1-I1347F (pink), (B) Na_V_1.1-I1347T (purple). And (C) Na_V_1.1-I1347V (blue).

**Supplemental Table S1.**
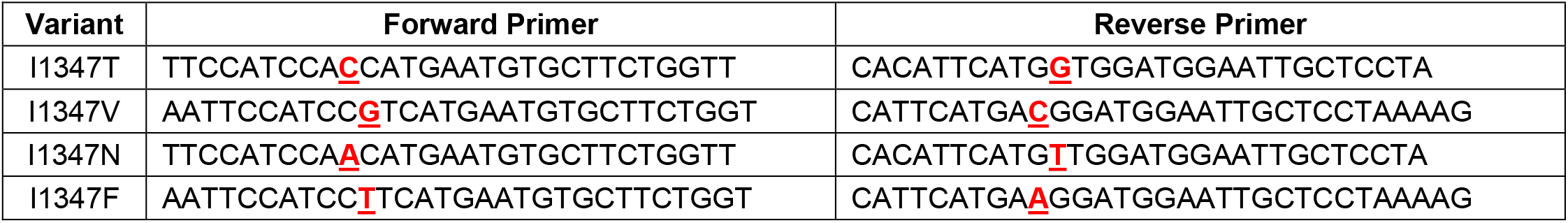
Mutagenic *SCN1A* primers.

**Supplemental Table S2.** Quantitative Data and Statistical Comparisons of *SCN1A* Variants (statistically significant values are in **bold**.)

| Peak Current Density<br>(0 mV, pA/pF) | WT 1 | I1347T | WT 2 | I1347V | I1347F |
| --- | --- | --- | --- | --- | --- |
| Mean ± SEM | -177.9 ± 29.1 | **-85.9 ± 18.5** | -171.9 ± 21.2 | **-109.6 ± 14.9** | **-108.7 ± 18.6** |
| n | 18 | **20** | 18 | **18** | **17** |
| p-value |  | **0.0099** |  | **0.0219** | **0.0329** |
| **V<sub>1/2</sub> Activation (mV)** |  |  |  |  |  |
| Mean ± SEM | -18.5 ± 1.6 | **-30.6 ± 1.2** | -18.3 ± 1.1 | -21.5 ± 1.6 | -19.4 ± 1.9 |
| n | 15 | **16** | 18 | 16 | 16 |
| p-value |  | **<0.0001** |  | 0.0987 | 0.6092 |
| **V<sub>1/2</sub> Inactivation (mV)** |  |  |  |  |  |
| Mean ± SEM | -60.0 ± 1.0 | **-67.0 ± 1.3** | -59.4 ± 1.6 | -63.0 ± 1.8 | -62.5 ± 1.7 |
| n | 15 | **16** | 18 | 17 | 17 |
| p-value |  | **0.0002** |  | 0.1382 | 0.2012 |
| **Recovery Tau Fast (ms)** |  |  |  |  |  |
| Mean ± SEM | 4.4 ± 0.4 | **11.6 ± 1.7** | 4.1 ± 0.6 | 5.2 ± 0.8 | 3.5 ± 0.5 |
| n | 12 | **11** | 13 | 16 | 16 |
| p-value |  | **0.0004** |  | 0.3075 | 0.4509 |
| **Recovery Tau Slow (ms)** |  |  |  |  |  |
| Mean ± SEM | 210.0 ± 15.7 | **374.5 ± 40.9** | 175.7 ± 28.0 | **280.4 ± 28.5** | 239.6 ± 42.1 |
| n | 13 | **13** | 13 | **16** | 16 |
| p-value |  | **0.0010** |  | **0.0154** | 0.2402 |
| **Recovery % Fast** |  |  |  |  |  |
| Mean ± SEM | 40.9 ± 2.7 | 35.1 ± 3.8 | 48.4 ± 2.5 | **40.9 ± 2.1** | **60.4 ± 2.7** |
| n | 13 | 13 | 15 | **16** | **16** |
| p-value |  | 0.2234 |  | **0.0291** | **0.0027** |
| **Use-Dependent Rundown**<br>**(P<sub>300</sub>/P<sub>1</sub>)** |  |  |  |  |  |
| Mean ± SEM | 77.4 ± 3.1 | **55.6 ± 6.4** | 81.5 ± 2.3 | **71.4 ± 3.7** | 81.4 ± 3.3 |
| n | 13 | **15** | 16 | **18** | 16 |
| p-value |  | **0.0072** |  | **0.0318** | 0.9757 |
| **Inactivation Tau at -30 mV (ms)** |  |  |  |  |  |
| Mean ± SEM | 2.3 ± 0.4 | 1.9 ± 0.2 | 3.4 ± 0.5 | 2.6 ± 0.4 | 4.8 ± 0.7 |
| n | 14 | 15 | 18 | 17 | 17 |
| p-value |  | 0.3262 |  | 0.4031 | 0.1098 |
| **Inactivation Tau at -20 mV (ms)** |  |  |  |  |  |
| Mean ± SEM | 1.5 ± 0.2 | 1.2 ± 0.1 | 1.5 ± 0.1 | 1.6 ± 0.2 | **2.2 ± 0.2** |
| n | 15 | 15 | 18 | 18 | **17** |
| p-value |  | 0.1019 |  | 0.7455 | **0.0195** |
| **Inactivation Tau at -10 mV (ms)** |  |  |  |  |  |
| Mean ± SEM | 0.80 ± 0.06 | 0.8 ± 0.1 | 0.7 ± 0.1 | 1.0 ± 0.1 | **1.1 ± 0.1** |
| n | 15 | 15 | 18 | 18 | **17** |
| p-value |  | 0.6144 |  | 0.1129 | **0.0037** |
| **Inactivation Tau at 0 mV (ms)** |  |  |  |  |  |
| Mean ± SEM | 0.59 ± 0.04 | 0.7 ± 0.1 | 0.61 ± 0.04 | **1.0 ± 0.1** | **0.9 ± 0.1** |
| n | 15 | 15 | 18 | **18** | **17** |
| p-value |  | 0.0734 |  | **0.0330** | **0.0121** |

**Supplemental Table S2.** Quantitative Data and Statistical Comparisons of *SCN1A* Variants – *continued*
| **Inactivation Tau at 10 mV (ms)** | **WT 1** | **I1347T** | **WT 2** | **I1347V** | **I1347F** |
| --- | --- | --- | --- | --- | --- |
| Mean ± SEM | 0.45 ± 0.03 | **0.6 ± 0.1** | 0.53 ± 0.04 | 0.9 ± 0.1 | **0.9 ± 0.1** |
| n | 14 | **15** | 18 | 18 | **16** |
| p-value |  | **0.0180** |  | 0.0679 | **0.0125** |
| **Inactivation Tau at 20 mV (ms)** |  |  |  |  |  |
| Mean ± SEM | 0.48 ± 0.03 | **0.6 ± 0.1** | 0.48 ± 0.04 | 0.8 ± 0.1 | **1.0 ± 0.2** |
| n | 15 | **14** | 17 | 18 | **17** |
| p-value |  | **0.0423** |  | 0.1478 | **0.0103** |
| **Ramp Current (pC/nA)** |  |  |  |  |  |
| Mean ± SEM | 2.1 ± 0.2 | **8.1 ± 1.2** | 2.8 ± 0.5 | **6.1 ± 1.2** | **12.9 ± 3.3** |
| n | 12 | **15** | 14 | **16** | **15** |
| p-value |  | **0.0001** |  | **0.0213** | **0.0074** |
| **Persistent Current (% Peak)** |  |  |  |  |  |
| Mean ± SEM | 0.5 ± 0.1 | **2.1 ± 0.4** | 1.0 ± 0.1 | **2.7 ± 0.7** | **5.4 ± 1.6** |
| n | 12 | **15** | 11 | **16** | **13** |
| p-value |  | **0.0047** |  | **0.0487** | **0.0173** |
| **Window Current (Area)** |  |  |  |  |  |
| Mean ± SEM | 0.43 ± 0.02 | **0.9 ± 0.1** | 0.47 ± 0.04 | **0.77 ± 0.09** | 0.46 ± 0.06 |
| n | 13 | **15** | 18 | **15** | 16 |
| p-value |  | **0.0026** |  | **0.0029** | 0.8884 |

**Supplemental Table S3.** Summary of observed functional effects of *SCN1A* variants in comparison to WT channels.

|  | **I1347T** | **I1347V** | **I1347F** |
| --- | --- | --- | --- |
| **Peak current** | smaller (LoF) | smaller (LoF) | smaller (LoF) |
| **Voltage-dependent Activation** | more negative (GoF) | WT-like | WT-like |
| **Voltage-dependent Inactivation** | more negative (LoF) | WT-like | WT-like |
| **Recovery from Inactivation** | slower (LoF) | slower (LoF) | faster (GoF) |
| **Inactivation Tau** | larger (GoF) | larger (GoF) | larger (GoF) |
| **Ramp current** | larger (GoF) | larger (GoF) | larger (GoF) |
| **Persistent current** | larger (GoF) | larger (GoF) | larger (GoF) |
| **Frequency-dependent rundown** | more rundown (LoF) | more rundown (LoF) | WT-like |
| **Window current** | larger (GoF) | larger (GoF) | WT-like |

**Movie S1** Dynamic morph of WT and I1347N structures.

Morph of AlphaFold 3 structures for WT and I1347N. The D4/S6 helix is colored orange in the foreground with the neighboring D3 helices shown in purple. The location of isoleucine-1347 and the I1347N substitution are illustrated as blue side chains.

## References

1. Lindy, A.S. et al. Diagnostic outcomes for genetic testing of 70 genes in 8565 patients with epilepsy and neurodevelopmental disorders. Epilepsia 59, 1062–1071 (2018)

2. Sheidley, B.R. et al. Genetic testing for the epilepsies: A systematic review. Epilepsia 63, 375–387 (2022)

3. D’Gama, A.M. et al. Evaluation of the feasibility, diagnostic yield, and clinical utility of rapid genome sequencing in infantile epilepsy (Gene-STEPS): an international, multicentre, pilot cohort study. Lancet Neurol. 22, 812–825 (2023)

4. Barcia, G. et al. Genetic etiologies with a large NGS panel in a monocentric cohort of 1000 patients with pediatric onset epilepsies. Epilepsia Open (2025, in press)

5. Scheffer, I.E. & Nabbout, R. SCN1A-related phenotypes: Epilepsy and beyond. Epilepsia 60 Suppl 3, S17–S24 (2019)

6. Mei, D., Cetica, V., Marini, C. & Guerrini, R. Dravet syndrome as part of the clinical and genetic spectrum of sodium channel epilepsies and encephalopathies. Epilepsia 60 Suppl 3, S2–S7 (2019)

7. Meisler, M.H., Hill, S.F. & Yu, W. Sodium channelopathies in neurodevelopmental disorders. Nat. Rev. Neurosci. 22, 152–166 (2021)

8. George, A.L. Jr. Emerging gene-based pathways: Channelopathies (epilepsy and beyond). In Swaiman’s Pediatric Neurology (7th ed., pp. 455–462). Elsevier, Philadelphia, PA (2025)

9. Catterall, W.A. Dravet syndrome: A sodium channel interneuronopathy. Curr. Opin. Physiol. 2, 42–50 (2018)

10. Bryson, A. & Petrou, S. SCN1A channelopathies: Navigating from genotype to neural circuit dysfunction. Front. Neurol. 14, 1173460 (2023)

11. Brunklaus, A. et al. The gain of function SCN1A disorder spectrum: novel epilepsy phenotypes and therapeutic implications. Brain 145, 3816–3831 (2022)

12. Wirrell, E.C. et al. International consensus on diagnosis and management of Dravet syndrome. Epilepsia 63, 1761–1777 (2022)

13. Richards, S. et al. Standards and guidelines for the interpretation of sequence variants: a joint consensus recommendation of the American College of Medical Genetics and Genomics and the Association for Molecular Pathology. Genet. Med. 17, 405–424 (2015)

14. Heyne, H.O. et al. Predicting functional effects of missense variants in voltage-gated sodium and calcium channels. Sci. Transl. Med. 12 (2020)

15. Brunklaus, A. et al. Gene variant effects across sodium channelopathies predict function and guide precision therapy. Brain 145, 4275–4286 (2022)

16. Brunklaus, A., George, A.L. Jr., Lal, D., Heinzen, E.L. & Goldman, A.M. Prophecy or empiricism? Clinical value of predicting versus determining genetic variant functions. Epilepsia 64, 2909–2913 (2023)

17. DeKeyser, J.M., Thompson, C.H. & George, A.L. Jr. Cryptic prokaryotic promoters explain instability of recombinant neuronal sodium channels in bacteria. J. Biol. Chem. 296, 100298 (2021)

18. Bochkov, Y.A. & Palmenberg, A.C. Translational efficiency of EMCV IRES in bicistronic vectors is dependent upon IRES sequence and gene location. Biotechniques 41, 283–292 (2006)

19. Thompson, C.H., Ben-Shalom, R., Bender, K.J. & George, A.L. Jr. Alternative splicing potentiates dysfunction of early onset epileptic encephalopathy SCN2A variants. J. Gen. Physiol. 152, e201912442 (2020)

20. Ganguly, S., Thompson, C.H. & George, A.L. Jr. Enhanced slow inactivation contributes to dysfunction of a recurrent SCN2A mutation associated with developmental and epileptic encephalopathy. J. Physiol. 599, 4375–4388 (2021)

21. Abramson, J. et al. Accurate structure prediction of biomolecular interactions with AlphaFold 3. Nature 630, 493–500 (2024)

22. Mariani, V., Biasini, M., Barbato, A. & Schwede, T. lDDT: a local super-position-free score for comparing protein structures and models using distance difference tests. Bioinformatics 29, 2722–2728 (2013)

23. de Kovel, C.G. et al. Targeted sequencing of 351 candidate genes for epileptic encephalopathy in a large cohort of patients. Mol. Genet. Genomic Med. 4, 568–580 (2016)

24. Landrum, M.J. et al. ClinVar: public archive of relationships among sequence variation and human phenotype. Nucleic Acids Res. 42, D980–D985 (2014)

25. Vaher, U. et al. De novo SCN8A mutation identified by whole-exome sequencing in a boy with neonatal epileptic encephalopathy, multiple congenital anomalies, and movement disorders. J. Child Neurol. 29, NP202–206 (2014)

26. Barker, B.S. et al. The SCN8A encephalopathy mutation p.Ile1327Val displays elevated sensitivity to the anticonvulsant phenytoin. Epilepsia 57, 1458–1466 (2016)

27. Johannesen, K.M. et al. Genotype-phenotype correlations in SCN8A-related disorders reveal prognostic and therapeutic implications. Brain 145, 2991–3009 (2022)

28. Vanoye, C.G. et al. High-throughput evaluation of epilepsy-associated KCNQ2 variants reveals functional and pharmacological heterogeneity. JCI Insight 7, e156314 (2022)

29. Thompson, C.H. et al. Epilepsy-associated SCN2A (NaV1.2) variants exhibit diverse and complex functional properties. J. Gen. Physiol. 155, e202313375 (2023)

30. Berg, A.T. et al. Expanded clinical phenotype spectrum correlates with variant function in SCN2A-related disorders. Brain 147, 2761–2774 (2024)

31. Vanoye, C.G. et al. Molecular and cellular context influences SCN8A variant function. JCI Insight 9 (2024)

32. Pablo, J.L.B. et al. Scanning mutagenesis of the voltage-gated sodium channel NaV1.2 using base editing. Cell Rep. 43, 114327 (2024)

33. Boßelmann, C.M., Hedrich, U.B.S., Lerche, H. & Pfeifer, N. Predicting functional effects of ion channel variants using new phenotypic machine learning methods. PLoS Comput. Biol. 19, e1010959 (2023)

